# Generation of parthenocarpic tomato plants in multiple elite cultivars using the CRISPR/Cas9 system

**DOI:** 10.1101/2023.04.27.538515

**Authors:** Cam Chau Nguyen, Tien Van Vu, Rahul Mahadev Shelake, Nhan Thi Nguyen, Tran Dang Khanh, Jae-Yean Kim

**Affiliations:** Division of Applied Life Science (BK21 Four Program), Plant Molecular Biology and Biotechnology Research Center, Gyeongsang National University, Jinju, Korea; Faculty of Biotechnology, Vietnam National University of Agriculture, Hanoi, Vietnam; Institute of Environmental Technology, Vietnam Academy of Science and Technology, Hanoi, Vietnam; Agricultural Genetics Institute, Hanoi, Vietnam

**Keywords:** IAA9, auxin signaling, CRISPR/Cas9, elite tomato, parthenocarpy, SlANT1

## Abstract

Tomato (*Solanum lycopersicum* L.) is one of the most important crops in the world for its fruit production. Advances in cutting-edge techniques have enabled the development of numerous critical traits related to the quality and quantity of tomatoes. Genetic engineering techniques, such as gene transformation and gene editing, have emerged as powerful tools for generating new plant varieties with superior traits. In this study, we induced parthenocarpic traits in a population of elite tomato (ET) lines. At first, the adaptability of ET lines to genetic transformation was evaluated to identify the best-performing lines by transforming the *SlANT1* gene overexpression cassette and then later used to produce the *SlIAA9* knockout lines using the CRISPR/Cas9 system. ET5 and ET8 emerged as excellent materials for these techniques and showed higher efficiency. Typical phenotypes of knockout *sliaa9* were clearly visible in G0 and G1 plants, in which simple leaves and parthenocarpic fruits were observed. The high efficiency of the CRISPR/Cas9 system in developing new tomato varieties with desired traits in a short period was demonstrated by generating T-DNA-free homozygous *sliaa9* knockout plants in the G1 generation. Additionally, a simple artificial fertilization method was successfully applied to recover seed production from parthenocarpic plants, securing the use of these varieties as breeding materials.

## Introduction

Tomato (*Solanum lycopersicum* L.) is an important fruit vegetable crop with diverse applications in food production and health benefits. The quality and quantity of tomato production has significantly improved in recent years (Sharma et al. 2019; Salava et al. 2021). The breeding, including conventional breeding and modern biotechnological tools, have been involved in producing elite tomato (ET) cultivars with superior traits (Abewoy Fentik 2017; Sharma et al. 2019; Xia et al. 2021). New plant breeding technologies, such as targeted mutagenesis, have been widely adopted to quickly introduce desired traits in tomatoes (Bai 2017; Vu et al. 2020a; Tiwari et al. 2023). Gene editing (GE) systems, including zinc-finger nucleases (ZFNs), transcription activator-like effector nucleases (TALENs), clustered regularly interspaced short palindromic repeats (CRISPR) with CRISPR-associated protein 9 (Cas9) (CRISPR/Cas9), base editors (BEs), and prime editors (PEs), have been applied in crops to target specific genes with high accuracy and produce transgene-free plants (Zhu & Zhu 2022).

Among these tools, CRISPR/Cas9 has shown significant potential in editing beneficial genes, promoting elite tomato production (Li et al. 2018; Pan et al. 2016; Reem & van Eck 2019; Santillán Martínez et al. 2020; Yang et al. 2022). This technology has been used to enhance biotic and abiotic stress resistance, improve fruit quality, and accelerate the domestication of tomatoes (Wang et al. 2019; Tiwari et al. 2023). However, the efficiency of GE tools depends on several factors, including the expression level of sgRNA and Cas proteins, the delivery efficiency of CRISPR/Cas reagents, the specificity of the gRNAs, and the transformation and regeneration systems (Ma et al. 2023).

Among the desired traits in tomatoes, parthenocarpic fruits are in high demand due to their better taste and higher industrial value (Gustafson 1942; Ueta et al. 2017; Xia et al. 2021). Modifying auxin signaling genes such as *SlIAA9, SlARF7*, and *SlARF5* resulted in parthenocarpic tomatoes, confirming the critical role of these phytohormones in seed production (Hu et al. 2018; Ueta et al. 2017). Knocking out the MADS-box gene *SlAGAMOUS-LIKE 6 (SlAGL6)* by CRISPR/Cas9 system resulted in facultative parthenocarpy and improved fruit yield under heat stress conditions (Klap et al. 2017). However, efficient modification of these parthenocarpy- related genes in commercial tomato cultivars is still limited. *SlAGL6* was successfully mutated in M82 cultivar to induce parthenocarpy (Klap et al. 2017), while seedless phenotype was observed only in Micro-Tom *sliaa9* edited lines (Saito et al. 2011), but not in M82 and Ailsa Craig (AC) backgrounds (Tran et al. 2021; Ueta et al. 2017). The *sliaa9* parthenocarpic breeding lines were generated by crossing gene-edited Micro-Tom and inbred tomato lines. Although the mutant allele was adopted, only 35.8% of fruits in the T_4_ population were seedless, and the fruit weight was significantly reduced (Tran et al. 2021). *Slagl6*/M82 (*sg*), on the other hand, produced around 80% of seedless fruits in the G1 population (Klap et al. 2017). These data raised the need to develop GE tools that are more efficient to direct introduce mutation to the parthenocarpy-related genes such as *SlIAA9* in the commercial tomato lines.

*SlIAA9*, a member of the Aux/IAA family, is the key regulator of auxin signaling related to parthenocarpy induction (Wang et al. 2005). This negative regulator of the auxin response pathway prevents ovary development before pollination by suppressing auxin signaling in flowers. The down-regulated mutants of *SlIAA9* released an auxin signal in young flowers, thus activating the auxin-responsive genes which initiate fruits without fertilization. The *sliaa9* mutant plants are not shown a reduction in fruit yield, fruit weight or flesh consistency (Mazzucato et al. 2015), suggesting a wonderful option to produce parthenocarpic lines while maintaining other superior traits of background cultivars. Besides the parthenocarpic phenotype, *sliaa9* mutants showed simple leave architecture, which is a convenient, visible marker for breeding and selection on a large scale. However, the performance of the *sliaa9* mutants differs depending on the genetic backgrounds and the cultivating conditions (Abe-Hara et al. 2021; Rahmat et al. 2023).

In recent years, many breeding programs resulted in many tomato cultivars with improved quantity and quality traits (Cappetta et al. 2020). Productivity, sweetness, fruit shape, ripening time, and biotic and abiotic stress tolerance are focused on by tomato breeders (Acquaah 2016). Recently, a collection of elite tomato (ET) lines was developed in Vietnam through conventional breeding associated with genetic markers selection (Nguyen et al. 2023). The F_8_ populations of ET lines showed superior traits such as significantly high yield, early maturing, jointless pedicels, partial parthenocarpy, and harboring *TYLCV*-resistant alleles. These newly developed tomato lines not only enable higher tomato production but also provide an excellent source material for future breeding programs.

To evaluate the efficacy of GE tools in the newly developed ET lines and produce parthenocarpic tomato lines that retain superior traits and adapt well to the local conditions of Vietnam, we employed CRISPR/Cas9-mediated GE system by establishing the tissue culture system for F8 plants and disrupt the function of *SlIAA9* gene. The transformation and GE efficiencies were evaluated in the G0 plants. The phenotypic traits of fruits and leaves were estimated in the G0 and G1 plants, revealing the variation in adopting genetic modification tools of different tomato lines. Finally, artificial fertilization was applied to recover seeds from the *SlIAA9*-edited lines to verify that the observed seedless phenotypes were derived from the true parthenocarpy.

## Materials and Methods

### Plant materials and growth conditions

Ten Vietnamese varieties (ET1 – ET10) which were agronomically examined in the previous study (Nguyen et al. 2023) were used as materials for genetic transformation and GE study. Hongkwang (HK) commercial variety was used as a control to evaluate the transformation efficiency and GE during all the experiments (Vu et al. 2020b). Transgenic plants were grown in soil pots at the greenhouse of the Gyeongsang National University of Agriculture (GNU) under optimized conditions.

### Plasmids and transformation

*pANT1ox* plasmid (Vu et al. 2020b) containing expression cassette of *SlANT1* driven by CAMV35S promoter was transformed into tested tomato lines to examine gene transformation efficiency.

In order to knockout the *SlIAA9* gene, the *pGE-IAA9* vector was cloned following MoClo (Weber et al. 2011) and Golden Gate (Engler et al. 2014) protocols to introduce indel mutation to exon 1 of *SlIAA9*. The construct contains a Cas9 expression cassette driven by the CaMV 2×35S promoter and 5’UTR and guide RNA (Ueta et al. 2017) scaffold driven by the AtU6 promoter (Kamoun Lab, Addgene #46968). The *AtU6-gRNA-7xT* fragment was cloned into level-1 plasmid (*SlIAA9-gRNA3*) (Supplementary. Fig. 1a). Enzyme digestion by *BpiI* of three clones shows the expected 4352 bp and 203 bp bands, and Sanger sequencing confirmed the correct level-1 sequences (Supplementary Fig. 1b-c). The level-2 construct *pGE-IAA9* was designed and cloned using Golden Gate system. The expression cassette of Kanamycin resistance gene (*NptII*) under the regulation of NOS promoter (*pNOS*) and OCS terminator (*tOCS*) with the omega enhancer sequence was placed in position 1 of *pGE-IAA9* (Fig. 2a). SpCas9 driven by 2×35S CaMV promoter and NOS terminator and *AtU6-9gRNA-7xT* were placed in position 2 and position 3 of the level-2 plasmid, respectively. Enzyme digestion using *NheI* and *PmlI* showed expected bands of 5333 bp, 3977 bp, and 2758 bp. Additionally, Sanger sequencing revealed that the construct was cloned successfully with correct junctions (Supplementary Fig. 2).

Seven-day-old cotyledons were used to prepare explants for *Agrobacterium*-mediated transformation (Vu et al. 2020b). *pANT1ox* and *pEG-IAA9* vectors were transformed into *Agrobacterium tumefaciens* GV3101 (pMP90) using the electroporation method. Ten and seven ET lines were used as materials for the transformation of *pANT1ox* and *pEG-IAA9*, respectively. HK cultivar was used as a control. Purple spots were counted on 21 days post-transformation (dpt). Transgenic shoots were grown in a growth chamber under long-day conditions (16 h light/8 h dark in 25°C) until roots were generated and transferred to soil pots. Transgenic plants were grown in the greenhouse until harvest. T-DNA insertion was determined by amplifying a 740-bp fragment within Kanamycin resistant gene located between the left border and right border of Ti plasmid using primer set (NptII-F5, 5’-TGGAGAGGCTATTCGGCTATG-3’ and NptII-R5, 5’-CTCGTCAAGAAGGCGATAGAA-3’). Internal control, which is Glyceraldehyde-3-phosphate dehydrogenase (GAPDH), was amplified by the primers GAPDH-F, 5’-GATTCGGAAGAATTGGCCG-3’ and GAPDH-R, 5’-TCATCATACACACGGTGAC-3’ to generate a 606 bp fragment.

### Tomato tissue culture

Explants transformed by *A. tumefaciens* were transferred to co-cultivation media containing ABM-MS and 200 μM acetosyringone and were kept under dark condition (25°C) for two days. Samples were washed with distilled water containing 500 μM timentin, then transferred to selection media SEL4 (MS Gamborg B5 vitamins 4.4 g/L, MES 0.976 g/L, Maltose 30 g/L, IAA 0.05 mg/L, Zeatin ribose trans isomer 0.5 mg/L, putrescine 1 mM, pH 5.7) containing 300 μM timentin and 80 μM kanamycin. Transformed explants were grown at 28°C for 5 days under short-day conditions (8 h light/16 h dark) and moved to 25°C growth chamber under long-day conditions (16 h light/8 h dark). Explant subculture to new media was performed every 14 days. The developed true shoots of explants were transferred to rooting media (MS salt containing B5 vitamins, sucrose 20 g/L, NAA 0.1 μM, IBA 0.3 μM, pH 5.8, timentin 300 μM). Transgenic plants were transferred to soil and grown in a greenhouse as described before (Vu et al. 2020b).

### Sanger sequencing and ICE analysis

Genomic DNA was collected from at least three different leaves of each plant. Edited alleles of *SlIAA9* were identified by conventional PCR from extracted genomic DNA of G0 and G1 leaves using primer set (IAA9-F1, 5’-CCCAGGATTGTAGGAGTTATGACATAC-3’ and IAA9-R1, 5’-GCCAACCAACAACCTGTGCCCTAA-3’) flanking region covering cutting site in exon 1. PCR products were separated in agarose gel 1%, then the specific bands were purified by BioFACT HiGeneTM Gel & PCR purification system (Cat. No. GP104-100). Sanger sequencing was performed by SolGent Analysis Service (SolGent Co., Korea) with the sequencing primer (IAA9-sF1, 5’-GAGCTGTGCAGGATTATTAGAG-3’). Edited alleles were visualized on SnapGene software and decomposed using ICE analysis (https://ice.synthego.com).

With samples that produced multiple alleles which could not be decomposed by ICE, PCR products were cloned to pJET1.2/blunt vectors using the CloneJET PCR Cloning kit (Thermo Scientific, Cat. Nos. K1231). Cloned plasmids were transformed into *E. coli* competent cells using the heat shock method, then the transformed cells were grown in Carbenicillin 250 mg/L media. Six colonies of each plate were analyzed by Sanger sequencing following plasmid extraction and PCR.

### Artificial fertilization

Blooming flowers were shaken with an electric brush for 5–10 seconds. Artificial pollination was performed repeatedly every day in the morning (9:00-10:00 am) during the flowering period.

## Results

### Transformation efficiency of F_8_ lines

In order to examine the ability to adopt foreign genes, the transformation of *pANT1ox* wa performed in all ten ET lines with HK as a control. Overexpression of *SlANT1* by *pANT1ox* construct (Fig. 1a) caused purple spots on tomato callus, in which each spot is correlated to one transformed cell (Fig. 1b). These purple cells can generate purple shoots and plants with purple leaves and fruits (Fig. 1b). After 21 days of transformation, purple spots were counted, and th results of three experiments were shown in Fig. 1c and Supplementary Table 1.

**Fig. 1.**
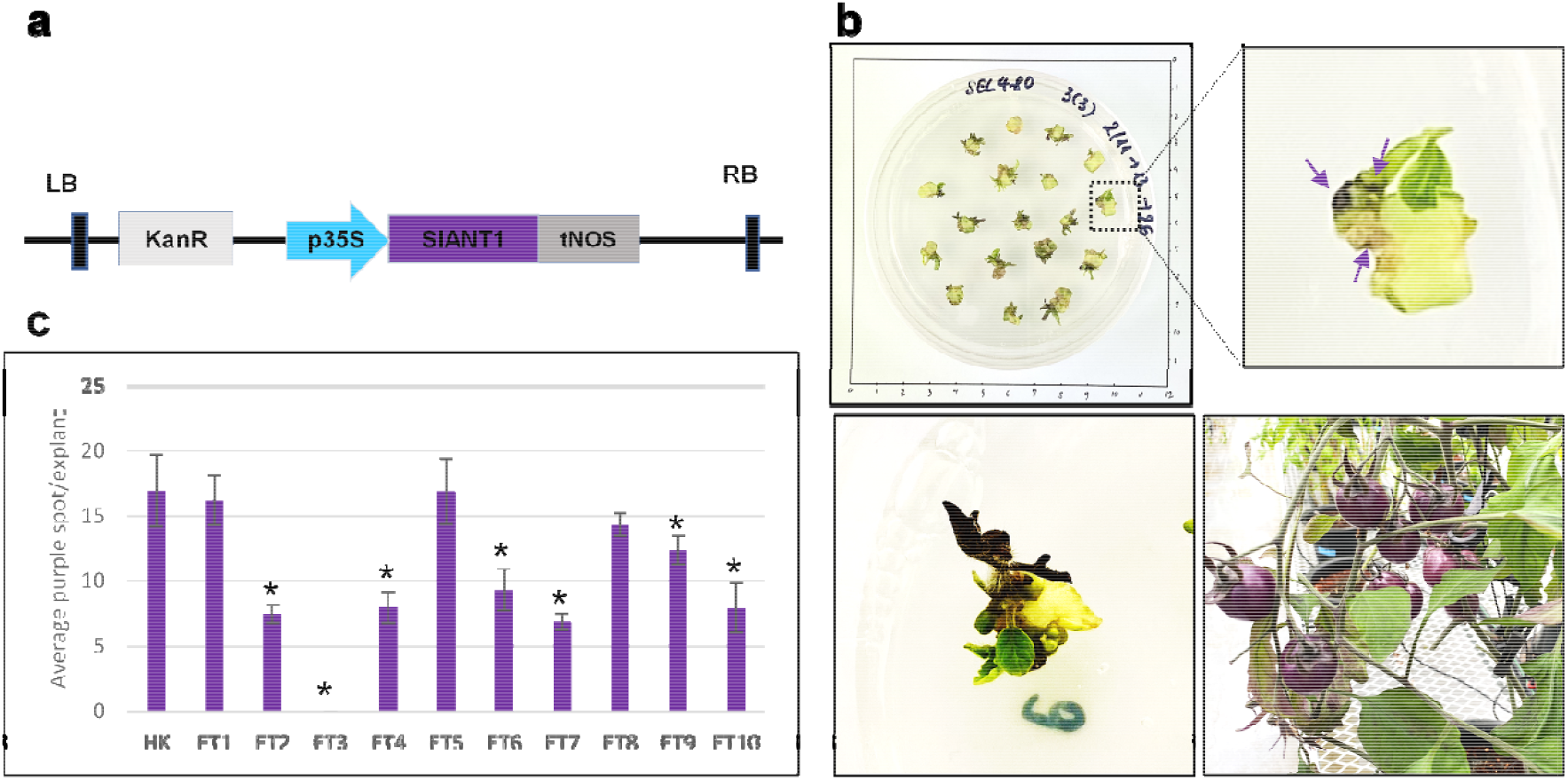
Genetic transformation efficiencies of elite tomato (ET) lines. 10 selected F8 lines were used as materials for transformation of *pANT1ox* construct. 7-day-old cotyledons were cut from seedlings and were transformed by *pANT1ox* constructs, then grown on Kanamycin selection media. At least 30 explants of each line were used in one experiment. Three biological repeats were conducted. **a** *pANT1ox* construct. KanR: kanamycin expression cassette (*pNOS-NptII-tOCS*), p35S: CAMV35S long-promoter. **b** *SlANT1* expression in tomato cotyledon (representative pictures). Upper panel: *SlANT1* expression at 21 days after transformation (DAT). Purple arrows indicated purple spots. Lower panel: purple shoots (left) and fruits (right). **c** Transformation efficiency. *y*-axis showed an average number of purple spots per explant; *x*-axis indicated tomato F8 lines tested in this experiment. Data represents the mean ± sd. n=3. An asterisk indicates a significant difference (*p* < 0.05) between ET lines and the control HK as determined by *t*-test.

HK cultivar has been widely used as a model commercial tomato variety in recent studies (Vu et al. 2020b; Tran et al. 2021) due to its higher genetic transformation efficiency and integration of T-DNA in the genome. Out of 10 lines tested, ET1 and ET5 exhibited transformation efficiencies comparable to the HK. While HK had the highest average number of purple spots (p.p.) (16.94 p.p./explant on average), ET1 and ET5 also showed an increased number of purple spots, which were 16.19 and 16.88 per explant on average, respectively. ET8 (14.32 p.p./explant) and ET9 (12.35 p.p./explant) displayed a slightly lower transformation efficiency than HK, but it was relatively high compared with the remaining lines. Remarkably, ET1 and ET8 were the progeny of the same parental lines, so their high transformation efficiency might be a heritable trait. However, this correlation was not observed between ET9 and ET4, which were generated from the same parents, suggesting segregation of the trait.

Notably, ET3 did not show any purple spot in all three replicates, indicating that *pANT1ox* was not expressed in this line (Fig. 1c and Supplementary Table 1). While the seedling of other lines exhibited purple shoots, ET3 had green shoots. Additionally, segregation lines of ET2, ET4, ET5, and ET9, which also had green shoots, were tested, but none of them showed any purple spots (Supplementary Table 2). These findings suggest that these lines are missing one or a few downstream factors of SlANT1. Based on the transformation efficiency and agronomical performance, 7 out of 10 ET lines were selected for GE experiments. Between ET1 and ET8, which were generated from the same parents’ combination, ET8 was chosen for being superior in productivity. ET3 and ET10 were eliminated due to low or undermined transformation efficiency (Fig. 1c).

### CRISPR/Cas9-based *SlIAA9* gene editing

To choose the most appropriate gRNA, the sequence of *SlIAA9* in the ten ET lines was compared with that of HK and Micro-Tom to examine the sequence variability. gRNA3 (20 bp) position showed 100% sequence match in the exon 1 of all the ET lines (Supplementary Fig. 3), so we chose this gRNA for knocking-out *SlIAA9* gene in the tested lines.

It was anticipated that the *pEG-IAA9* construct (Fig. 2a) would introduce indel mutations into exon 1 of the *SlIAA9* gene, thereby knocking this gene out or down in the ET lines used for research. The highest number of true shoots (63) and G0 plants (59) were generated from 120 transformed explants of ET8 (Table 1). Among them, 40 plants carried T-DNA insertion of th construct, which was 33.33%, and 19 plants, equal to 15.83% exhibited the simple leaves phenotype. Despite only 27 true shoots (27%) generated from ET5, we obtained 25 G0 plants, with 13% exhibiting the simple leaves phenotype, making ET5 the second most promising candidate for *SlIAA9* editing. The percentage of simple-leaf plants generated from ET2, ET4, and ET9 was relatively low at 3%, 2.5%, and 1.67%, respectively. ET6 and ET7 produced a small number of true shoots (10% and 11.25%), and no plants with the simple leaf phenotype were observed.

**Fig 2.**
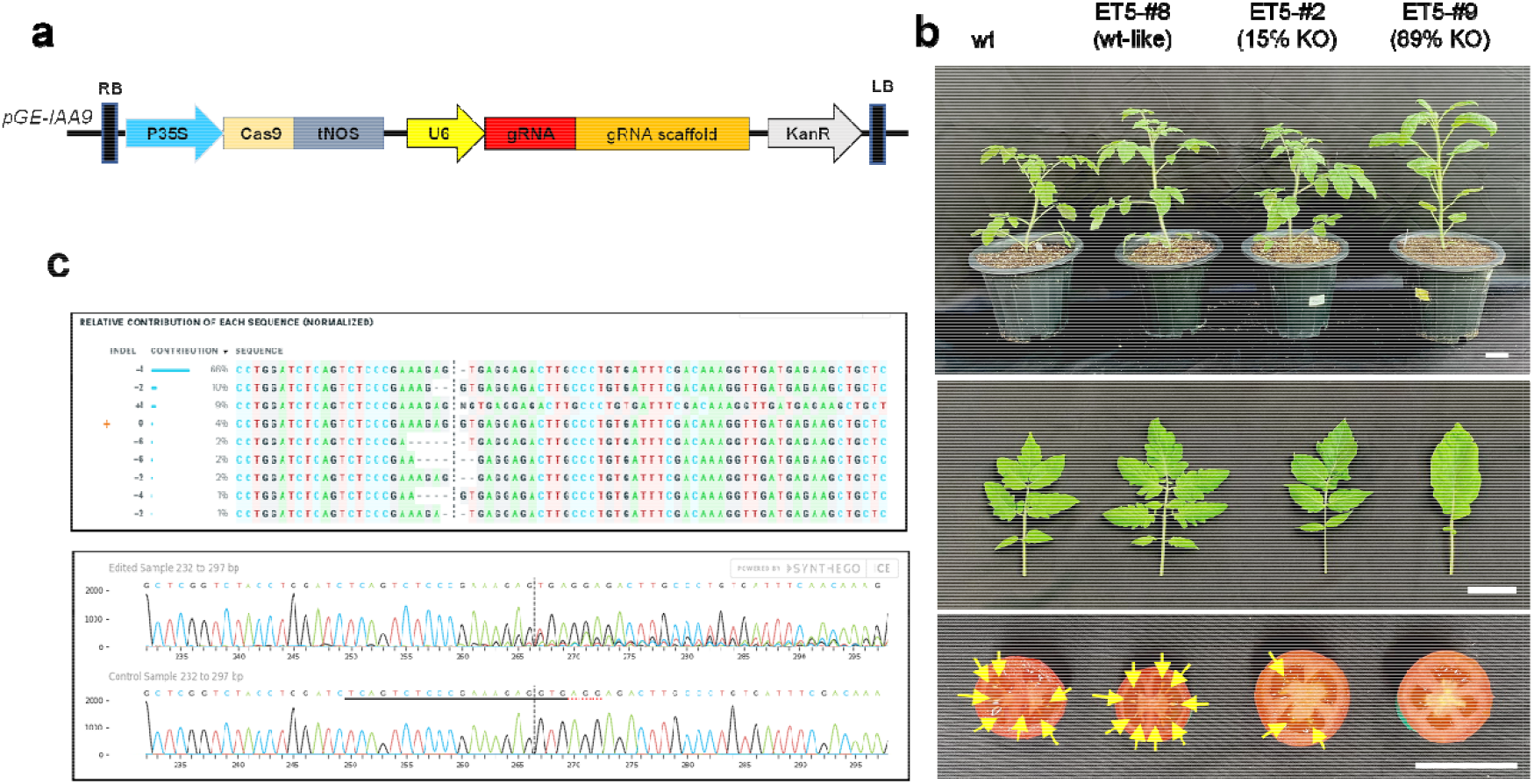
Editing of *SlIAA9* in elite tomato (ET) lines. Cotyledons of 7-day-old plants were transformed by Agrobacterium carrying *pGE-IAA9* plasmids, then grown and selected by antibiotic-containing shoot induction media. **a** Map of gene editing construct. Cas9 protein was expressed under the CAMV35S promoter, while sgRNA was expressed under AtU6 promoter. KanR: kanamycin expression cassette (*pNOS-NptII-tOCS*). **b** *SlIAA9*-edited G0 plants (representative pictures). Shoots were transferred into root induction media for 3 weeks before moving to the soil. Pictures of G0 plants (upper panel) and leaves (middle panel) were captured 4 weeks after moving to the soil. Yellow arrows in the lower panel indicate seeds in fruits of G0 individuals. Bar = 5cm. **c** Allele contribution in plant ET5-#9. *SlIAA9* sequences were analyzed by ICE-Synthego online software (https://ice.synthego.com). Sequences of mutated alleles were compared with wild-type (marked with yellow “+” symbol) and proportions of each allele were shown. The black vertical dotted line indicates the cut site.

**Table 1.**
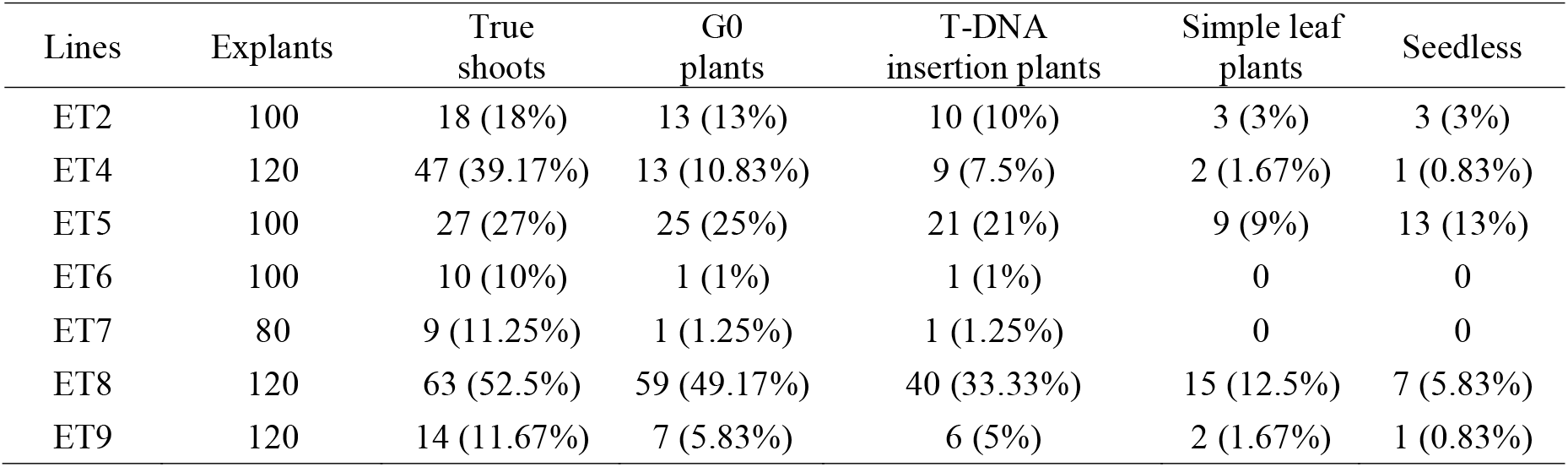
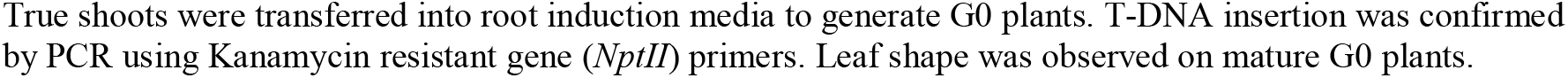
Assessment of transformants obtained from the transformation of pEG-IAA9 construct on tested tomato lines.

The leaf and fruit phenotypes of G0 plants were affected by the ratio of knockout alleles. For example, plant ET5-#2, which showed normal leaves, had 15% knockout alleles, whereas ET5-#9 had 89% knockout alleles and displayed simple leaf phenotypes (Fig. 2b). Mildly affected phenotype plants such as ET5-#2 fruits had few seeds, which was less than ten seeds per fruit (Fig. 2b). Severe knockout plants such as ET5-#9 showed parthenocarpic fruits. Up to 10 plants of each background were analyzed for allele contribution, mutant rate and knock out score using Sanger sequencing and pJET cloning (Table 2). All the G0 plants exhibiting the phenotype of simple leaves showed the presence of T-DNA.

**Table 2.** Genotyping and phenotyping of *SlIAA9* mutant G0 plants generated by CRISPR/Cas9 system.

| No. | Plant # | T-DNA insertion | Reading score | Mutation rate | KO score | Simple leaf | Seed phenotype |
| --- | --- | --- | --- | --- | --- | --- | --- |
| 1 | ET2-#1 | X | 1 | 0 | 0 | N | N |
| 2 | ET2-#2 | O | 0.97 | 0.52 | 0.52 | N | X, F |
| 3 | ET2-#3 | O | 0.94 | 0.94 | 0.47 | N | X |
| 4 | ET2-#4 | O | 0.94 | 0.94 | 0.48 | O | F |
| 5 | ET2-#5 | O | 0.98 | 0.27 | 0.27 | N | F |
| 6 | ET2-#6 | O | 0.97 | 0.2 | 0 | N | N |
| 7 | ET2-#7 | O | 1 | 0 | 0 | N | F |
| 8 | ET2-#8 | O | 0.9 | 0.62 | 0.62 | O | X |
| 9 | ET2-#9 | O | 1 | 0 | 0 | N | F |
| 10 | ET2-#10 | O | - | 0.33* | 0.16* | N | F |
| 11 | ET4-#1 | O | 1 | 0 | 0 | N | N |
| 12 | ET4-#2 | X | 1 | 0 | 0 | N | N |
| 13 | ET4-#3 | X | 1 | 0 | 0 | N | N |
| 14 | ET4-#4 | X | 1 | 0 | 0 | N | N |
| 15 | ET4-#5 | O | 1 | 0 | 0 | N | N |
| 16 | ET4-#6 | O | 1 | 0 | 0 | N | N |
| 17 | ET4-#7 | O | 0.97 | 0.49 | 0.49 | O | N |
| 18 | ET4-#8 | O | 1 | 0 | 0 | N | N |
| 19 | ET4-#9 | O | 1 | 0 | 0 | N | N |
| 20 | ET4-#10 | O | 0.96 | 0.67 | 0.67 | O | X |
| 21 | ET5-#1 | O | - | 1* | 1* | O | X |
| 22 | ET5-#2 | O | 0.86 | 0.17 | 0.17 | N | F |
| 23 | ET5-#3 | X | 1 | 0 | 0 | N | F |
| 24 | ET5-#4 | O | - | 0.83* | 0.83* | N | X |
| 25 | ET5-#5 | O | - | 1* | 0.83* | O | X |
| 26 | ET5-#6 | O | 0.96 | 0.68 | 0.08 | N | F |
| 27 | ET5-#7 | O | 0.97 | 0.93 | 0.93 | O | X |
| 28 | ET5-#8 | O | 1 | 0 | 0 | N | X |
| 29 | ET5-#9 | O | 0.97 | 0.93 | 0.89 | N, O | X |
| 30 | ET5-#10 | X | - | 0* | 0* | N | N |
| 31 | ET6-#1 | O | 0.98 | 0.2 | 0.14 | N | N |
| 32 | ET7-#1 | O | 0.91 | 0.91 | 0.61 | N | X |
| 33 | ET8-#1 | O | 0.97 | 0.93 | 0.93 | O | N |
| 34 | ET8-#2 | O | 1 | 0 | 0 | N | N |
| 35 | ET8-#3 | X | 1 | 0 | 0 | N | N |
| 36 | ET8-#4 | O | 0.99 | 0.19 | 0.19 | N | N |
| 37 | ET8-#5 | O | 0.98 | 0.94 | 0.94 | N, O | X |
| 38 | ET8-#6 | O | 0.96 | 0.51 | 0.51 | N | F |
| 39 | ET8-#7 | O | 0.96 | 0.95 | 0.85 | O | X, N |
| 40 | ET8-#8 | X | 1 | 0 | 0 | N | N |
| 41 | ET8-#9 | O | 0.97 | 0.85 | 0.8 | O | F |
| 42 | ET8-#10 | O | 0.97 | 0.8 | 0.71 | O | X |
| 43 | ET9-#1 | O | 0.99 | 0.07 | 0.07 | N | F |
| 44 | ET9-#2 | O | 0.95 | 0.56 | 0.43 | N | N |
| 45 | ET9-#3 | X | 1 | 0 | 0 | N | N |
| 46 | ET9-#4 | O | 1 | 0 | 0 | N | N |
| 47 | ET9-#5 | O | 1 | 0 | 0 | N | N |
| 48 | ET9-#6 | O | - | 1* | 1* | O | X |
| 49 | ET9-#7 | O | 0.95 | 0.95 | 0.95 | O | N |

KO, knockout; X, no T-DNA insertion/seedless; O, has T-DNA insertion/simple leave phenotype; N, normal leaf shape/ normal seed number; F, few seeds (<10 seeds/fruit). Asterisks (\*) represent data obtained by PJET cloning followed by Sanger sequencing and ICE analysis.

Sanger sequencing analysis followed by ICE analysis revealed that multiple alleles were generated in each edited G0 plant (Fig. 2c, Supplementary Fig. 4). Most frame-shift mutations led to truncated proteins (Supplementary Table 3). The phenotypes of the plant were determined by the dominant alleles in the pool. In the case of ET5-#9, 89% of alleles were knocked-out; and among them, 66% were one-nucleotide deletion alleles. This mixture of alleles resulted in simple leaves and seedless fruits phenotype. At G1 generation, homozygous plants of alleles such as -1, -2, -4, -5, -6, -7, +1 bp were generated. Heterozygous and chimeric genotypes were also observed and summarized in Table 3. In one case, a rather long deletion of 16 nucleotides at the cutting site of *SlIAA9* was inherited in ET2-#5-5. Additionally, ET8#10-1 showed a significantly long deletion allele of -218 bp, together with a -1 bp mutant allele, and both plants expressed simple leaves phenotype. Besides, all the homozygous knockout G1 plants showed a simple leaf phenotype (Fig. 3a, Table 3). Furthermore, there are heterozygous and chimeric plants that contain up to four edited alleles of *SlIAA9*. Indeed, four edited G1 plants were T-DNA-free, including a homozygous plant with two nucleotide deletion alleles (ET2-#2-5). These results indicated that the edited alleles were stably inherited in subsequent generations.

**Table 3.**
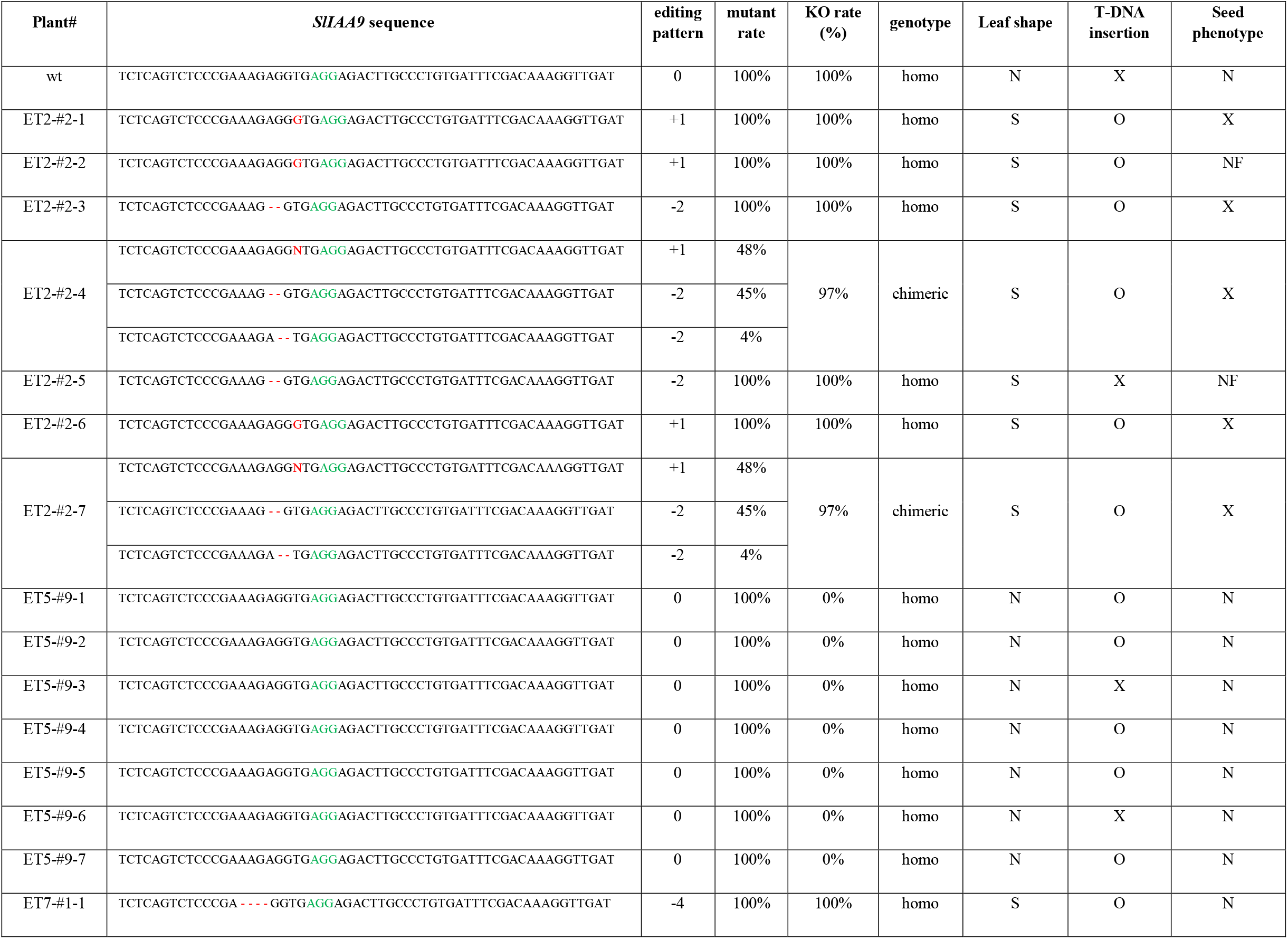
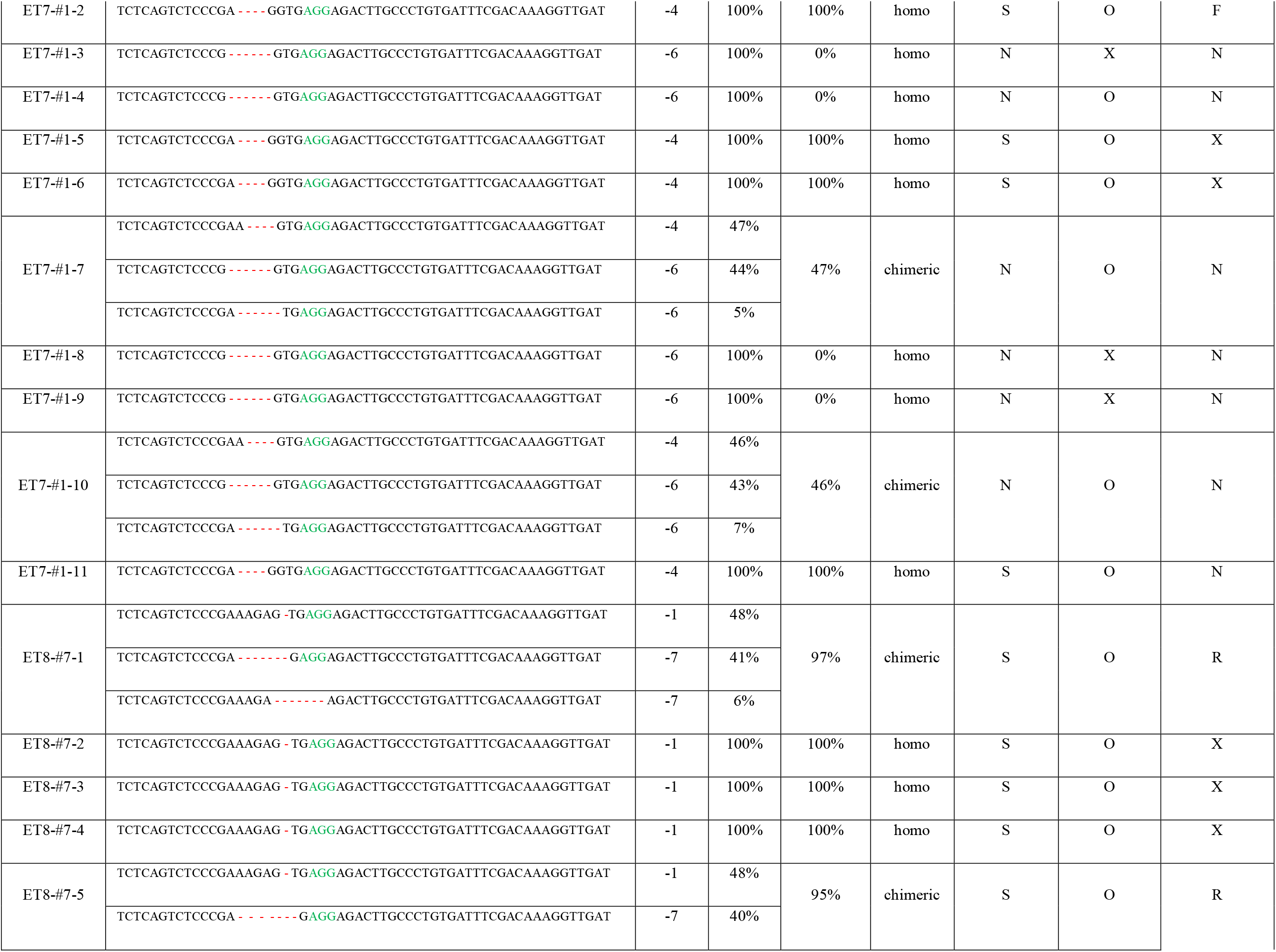
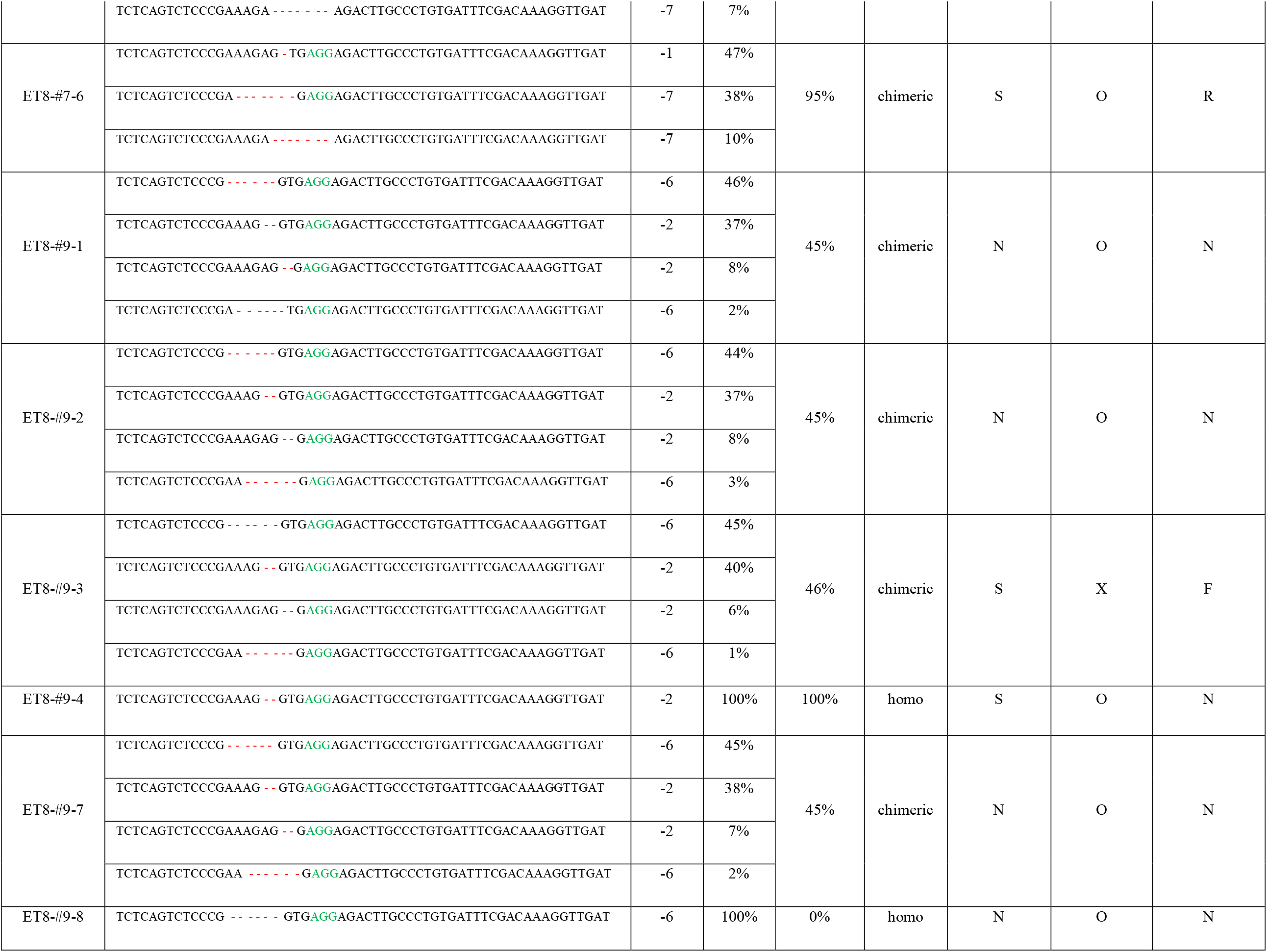
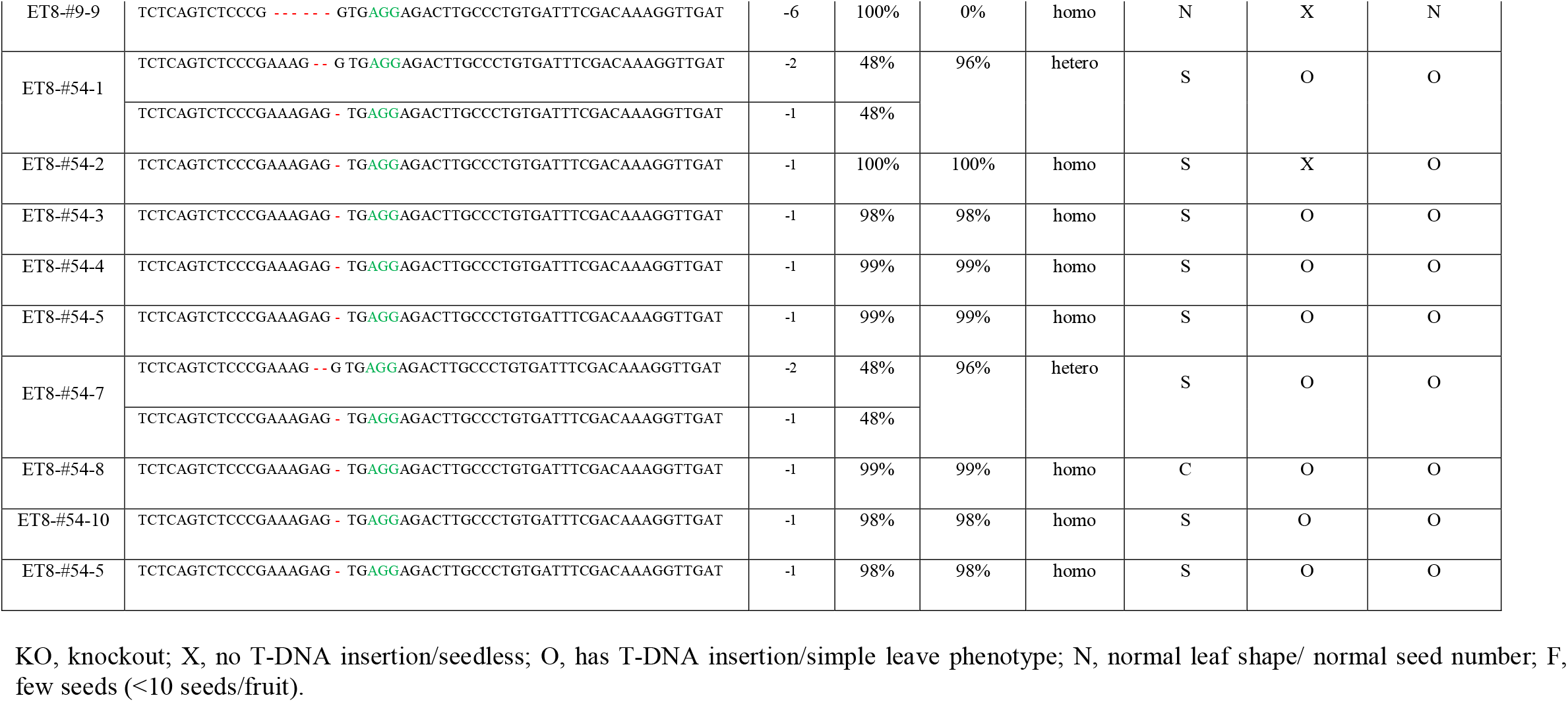
Genotyping and phenotyping of *sliaa9*-G1 generation.

**Fig 3.**
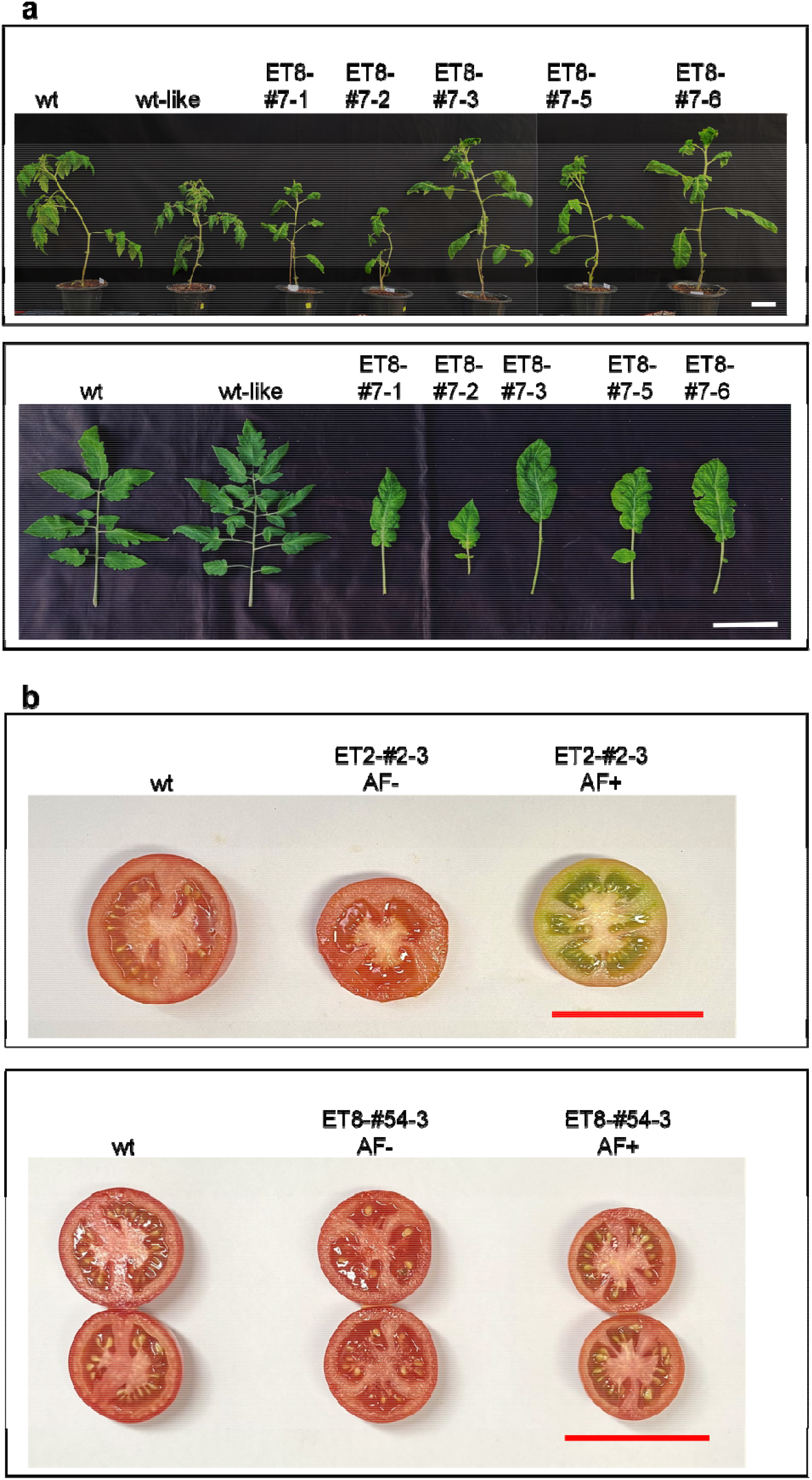
Phenotype of *SlIAA9*-knockout G1 generation. Seeds were obtained from G0 plants and were selected based on leaf phenotypes and T-DNA insertion. **a** Plant and leaf shape of representative *SlIAA9* editing G1 lines generated from ET8-#7 (G0). **b** Section phenotypes of G1 fruits of ET2-#2-3 (100% KO) and ET8-#54-3 (98% KO). wt, wild-type; AF-, fruits without artificial fertilization; AF+, Fruit with artificial fertilization. White bars = 10cm; red bars = 5cm.

### Artificial fertilization helps recover seeds from parthenocarpic lines

Artificial fertilization was conducted by shaking young to blooming flowers using an electric toothbrush. In the absence of artificial pollination, the original fruits of G1 knockout exhibited either no seed (ET2-#2-3) or a low number of seeds (ET8-#54-3). However, the artificially fertilized ones showed as many seeds as the wild types (Fig. 3b). This phenomenon was observed in various lines from different backgrounds, indicating that malfunctioning of *SlIAA9* promoted fruiting without fertilization, while pollen viability remained unaffected. The parthenocarpic phenotype, therefore, is greatly influenced by culturing conditions. This observation can serve as a useful method to obtain seeds for maintaining true parthenocarpic lines without the need for tissue culture or complicated propagation methods.

## Discussion

### Anthocyanin markers work selectively between the ET sublines

Anthocyanin production has been widely used as a visual selection marker in plant transformation as well as in CRISPR technology (Kortstee et al. 2011; Ruan et al. 2021). Overproduction of the *SlANT1* gene generated high anthocyanin pigment in transformants indicating the approximate level of transformation in plant tissues (Vu et al. 2020b). However, the result in Fig. 1c and Supplementary Table 1 suggest that the marker did not work in ET3, whose young shoot color was green instead of purple. A rough test experiment revealed that all the green shoot sub-lines of other ET lines failed to produce anthocyanin pigments despite the *pANT1ox* transformation (Supplementary Table 2). It is suggested that green-shoot tomato lines such as ET3 are missing one or a few downstream factors of SlANT1. One or more of the structural genes involved in SlANT1-dependent anthocyanin biosynthetic pathways might malfunction, including early enzymes such as CHS, CHI, and F3H, or late enzymes such as DFR and 3-GT (Mathews et al. 2003; Schreiber et al. 2012). These data proved that not all visual markers could be used to assist genome editing, and it is vital to understand the genetic background of the materials.

### Transformation and editing efficiency vary among tomato lines

The editing efficiency of *SlIAA9* differs between Micro-Tom and the commercial line AC (Ueta et al. 2017). The use of pEgP237-gRNA2(20bp)-2A-GFP resulted in a 100% mutation rate in Micro-Tom shoots but only 42.9% in AC shoots. Unlike Micro-Tom, AC edited plants were not reported to produce parthenocarpic fruits (Tran et al. 2021; Ueta et al. 2017). Table 1 and Table 2 reveal a high degree of variation in both transformation and editing efficiency among the tested population of ET lines. While some of these lines share parental origins, they do not exhibit the same level of foreign construct adoption potential. Notably, the transformation efficiency using *pANT1ox* plasmid and pEG-IAA9 are closely correlated in elite lines such as ET5 and ET8 (Table 1 and Supplementary Table 1). ET5 presented 16.88 purple spots per explant on average, and 21% of pEG-IAA9 transformed plants possessed T-DNA insertion. These numbers in ET8 were 14.32 and 33.33%, respectively. These two lines are the best F8 ET lines to be used as materials for genetic modification technologies due to their well-response to foreign gene transformation. Between these two lines, ET5 showed higher editing efficiency, presented by the amount of simple leaf and seedless plants in the G0 population (Table 1). However, the high productivity and survival rate of ET8 benefit this line to maintain and transfer the edited alleles to the next generations (Table 3). For the generation of commercial genome editing tomato, ET8 is the best-recommended option which provides beneficial traits such as high yield, high transformation efficiency, and low fruit cracking ratio (Nguyen et al. 2023).

### Seed number of parthenocarpic lines is culture-dependent

The number of seeds in *SlIAA9*-edited tomatoes exhibited variability not only between different editing events but also within each G1 subline that has a similar genetic background. In G0 and G1 plants, the observed phenotypes for seed number included the typical seed forming, a few seeds (less than ten seeds in each fruit), and zero seeds, which are known as seedless plants. Additionally, even within a single plant, each fruit could produce a different number of seeds. However, applying artificial fertilization to parthenocarpic plants significantly recovers seeds in G1 plants (Fig. 3b). This result revealed that culturing conditions such as physical impacts on blooming flowers, would quickly lead to a change in the seed number of the fruits. Parthenocarpy generated by modifying auxin biosynthesis and regulation pathways is known to induce fruit formation without fertilization (Wang et al. 2005; Wang et al. 2009), so the fruit formation would be initiated earlier than normal. It is predicted that natural physical impacts on young and blooming flowers, such as wind and insects, can restore seed formation in parthenocarpic fruits. This characteristic could be exploited to develop an efficient method for maintaining parthenocarpic lines and to consult the optimal conditions to produce a high proportion of seedless tomatoes.

## Author contribution

C.C.N. and J.-Y.K. conceived the idea and designed the experiments. C.C.N. designed experiments, performed experiments, analyzed data, and wrote the manuscript; C.C.N., T.V.V., and N.N.T. performed the experiments; R.M.S. contributed to the interpretation and draft of the output; R.M.S., J.-Y.K. and T.D.K revised the manuscript. All authors read and approved the final manuscript.

## Data availability

The data used in this study are available from the corresponding author upon reasonable request.

## Declarations

Not applicable.

## Ethics approval and consent to participate

Not applicable.

## Consent for publication

Not applicable.

## References

Abe-Hara C, Yamada K, Wada N, et al (2021) Effects of the sliaa9 mutation on shoot elongation growth of tomato cultivars. Front Plant Sci 12 :627832.

Abewoy Fentik D (2017) Review on genetics and breeding of tomato (Lycopersicon esculentum Mill). Advances in Crop Science and Technology 05:306.

Acquaah G (2015) Conventional plant breeding principles and techniques. In: Al-Khayri JM, Jain SM, Johnson DV (eds) Advances in plant breeding strategies: breeding, biotechnology and molecular tools. Springer International Publishing, Cham, pp 115–158.

Bai Y (2017) Developments in tomato breeding: conventional and biotechnology tools. In: Mattoo A, Handa A (eds) Achieving sustainable cultivation of tomatoes. Burleigh Dodds Science Publishing, UK.

Cappetta E, Andolfo G, Di Matteo A, et al (2020) Accelerating tomato breeding by exploiting genomic selection approaches. Plants 9:1236.

Engler C, Youles M, Gruetzner R, et al (2014) A Golden Gate modular cloning toolbox for plants. ACS Synth Biol 3: 839–843.

Gustafson FG (1942) Parthenocarpy: Natural and artificial. Bot. Rev.8: 599–654.

Hu J, Israeli A, Ori N, Sun TP (2018) The interaction between DELLA and ARF/IAA mediates crosstalk between gibberellin and auxin signaling to control fruit initiation in tomato. Plant Cell 30:1710–1728.

Klap C, Yeshayahou E, Bolger AM, et al (2017) Tomato facultative parthenocarpy results from SlAGAMOUS-LIKE 6 loss of function. Plant Biotechnol J 15:634–647.

Kortstee AJ, Khan SA, Helderman C, et al (2011) Anthocyanin production as a potential visual selection marker during plant transformation. Transgenic Res 20:1253–1264.

Mathews H, Clendennen SK, Caldwell CG, et al (2003) Activation tagging in tomato identifies a transcriptional regulator of anthocyanin biosynthesis, modification, and transport. Plant Cell 15:1689–1703.

Mazzucato A, Cellini F, Bouzayen M, et al (2015) A TILLING allele of the tomato Aux/IAA9 gene offers new insights into fruit set mechanisms and perspectives for breeding seedless tomatoes. Molecular Breeding 35:1–15.

Nguyen C-C, Shelake RM, Vu VT, et al (2023) Characterization of yield and fruit quality parameters of Vietnamese elite tomato lines generated through phenotypic selection and conventional breeding method. bioRxiv.

Rahmat BPN, Octavianis G, Budiarto R, et al (2023) SlIAA9 mutation maintains photosynthetic capabilities under heat-stress conditions. Plants 12:378.

Ruan Y, Chen K, Su Y, et al (2021) A root tip-specific expressing anthocyanin marker for direct identification of transgenic tissues by the naked eye in symbiotic studies. Plants 10:605.

Saito T, Ariizumi T, Okabe Y, et al (2011) TOMATOMA: A novel tomato mutant database distributing micro-tom mutant collections. Plant Cell Physiol 52: 283–296.

Salava H, Thula S, Mohan V, et al (2021) Application of genome editing in tomato breeding: Mechanisms, advances, and prospects. Int J Mol Sci 22:682.

Schreiber G, Reuveni M, Evenor D, et al (2012) ANTHOCYANIN1 from Solanum chilense is more efficient in accumulating anthocyanin metabolites than its Solanum lycopersicum counterpart in association with the ANTHOCYANIN FRUIT phenotype of tomato. Theor Appl Genet 124:295–307.

Sharma P, Thakur S, Negi R (2019) Recent advances in breeding of tomato-a review. Intl J Curr Microbiol App Sci 8:1275–1283.

Tran LT, Nguyen AT, Nguyen MH, et al (2021) Developing new parthenocarpic tomato breeding lines carrying iaa9-3 mutation. Euphytica 217:139.

Ueta R, Abe C, Watanabe T, et al (2017) Rapid breeding of parthenocarpic tomato plants using CRISPR/Cas9. Sci Rep 7:507.

Vu T Van, Das S, Tran MT, et al (2020a) Precision genome engineering for the breeding of tomatoes: recent progress and future perspectives. Front Genome Ed 2: 612137.

Vu T Van, Sivankalyani V, Kim E-J, et al (2020b) Highly efficient homology-directed repair using CRISPR/Cpf1-geminiviral replicon in tomato. Plant Biotechnol J 18:2133–2143.

Wang H, Jones B, Li Z, et al (2005) The tomato Aux/IAA transcription factor IAA9 is involved in fruit development and leaf morphogenesis. Plant Cell 17:2676–2692.

Wang H, Schauer N, Usadel B, et al (2009) Regulatory features underlying pollination-dependent and independent tomato fruit set revealed by transcript and primary metabolite profiling. Plant Cell 21:1428–1452.

Weber E, Engler C, Gruetzner R, et al (2011) A modular cloning system for standardized assembly of multigene constructs. PLoS One 6:e16765.

Xia X, Cheng X, Li R, et al (2021) Advances in application of genome editing in tomato and recent development of genome editing technology. Theor Appl Genet 134:2727–2747.

