## Supplementary Table 1, Supplementary Table 2, Supplementary Table 3, Supplementary Fig. 1, Supplementary Fig. 2, Supplementary Fig. 3 for "Generation of parthenocarpic tomato plants in multiple elite cultivars using the CRISPR/Cas9 system"

Supplementary Table 1. *pANT1ox* transformation efficiency of elite tomato lines

| Lines | Purple spots/explant | SD |
|---|---|---|
| HK | 16.94 | 2.74 |
| ET1 | 16.19 | 1.88 |
| ET2 | 7.48 | 0.70 |
| ET3 | 0.00 | 0.00 |
| ET4 | 7.98 | 1.20 |
| ET5 | 16.88 | 2.52 |
| ET6 | 9.32 | 1.57 |
| ET7 | 6.88 | 0.64 |
| ET8 | 14.32 | 0.88 |
| ET9 | 12.35 | 1.08 |
| ET10 | 7.93 | 1.87 |

Supplementary Table 2. *pANT1ox* expression in ET sub-lines

| Lines/sub-lines | Explant number | Purple spots | Purple spots/explant |
|---|---|---|---|
| ET2-gs | 27 | 0 | 0.00 |
| ET2 | 20 | 154 | 7.70 |
| ET3 | 27 | 0 | 0.00 |
| ET4-gs | 19 | 0 | 0.00 |
| ET4 | 18 | 116 | 6.44 |
| ET5-gs | 45 | 0 | 0.00 |
| ET5 | 47 | 769 | 16.36 |
| ET7-gs | 26 | 0 | 0.00 |
| ET7 | 17 | 102 | 6.00 |
| ET9-gs | 27 | 2 | 0.07 |
| ET9 | 24 | 251 | 10.46 |
| HK | 9 | 140 | 15.56 |

gs: Green-shoot sublines of ET lines

Supplementary Table 3. High frequent mutant alleles in G1 population of *SlIAA9*-edited plants

| Indel | Protein length | Mutant pattern | Sequences |
|---|---|---|---|
| 0 | 349 | wt | MSPPLLGVGEEEGQSNVTLLASSTSLGSICIKGSALKERNYMGLSDCSSVDSCNISTSSEDNNGCGLNLKATELRLGL PGSQSPERGEETCPVISTKVDEKLLFPLHPSKDTAFSVSQKTVVSGNKRGFSDAMDGFSEGKFLSNSGVKAGDTKET SRVQPPKMKDANTQSTVPERPSAVNDASNRAGSGAPATKAQVVGWPPIRSFRKNTLASASKNNEEVDGKAGSPAL FIKVSMDGAPYLRKVDLRTCSAYQELSSALEKMFSCFTIGQYGSHGAPGKDMLSESKLKDLLHGSEYVLTYEDKDG DWMLVGDVPWEMFIDTCKRLRIMKGSDAIGLPQGLWKSVGAEI* |
| -1 | 92 | truncated | MSPPLLGVGEEEGQSNVTLLASSTSLGSICIKGSALKERNYMGLSDCSSVDSCNISTSSEDNNGCGLNLKATELRLGL PGSQSPERVRRLAL* |
| -2 | 86 | truncated | MSPPLLGVGEEEGQSNVTLLASSTSLGSICIKGSALKERNYMGLSDCSSVDSCNISTSSEDNNGCGLNLKATELRLGL PGSQSPER* |
| -7 | 92 | truncated | MSPPLLGVGEEEGQSNVTLLASSTSLGSICIKGSALKERNYMGLSDCSSVDSCNISTSSEDNNGCGLNLKATELRLGL PGSQSPERVRRLAL* |
| -5 | 96 | truncated | MSPPLLGVGEEEGQSNVTLLASSTSLGSICIKGSALKERNYMGLSDCSSVDSCNISTSSEDNNGCGLNLKATELRLGL PGSQSPERGDLPCDFDKG* |
| +1 | 87 | truncated | MSPPLLGVGEEEGQSNVTLLASSTSLGSICIKGSALKERNYMGLSDCSSVDSCNISTSSEDNNGCGLNLKATELRLGL PGSQSPER* |

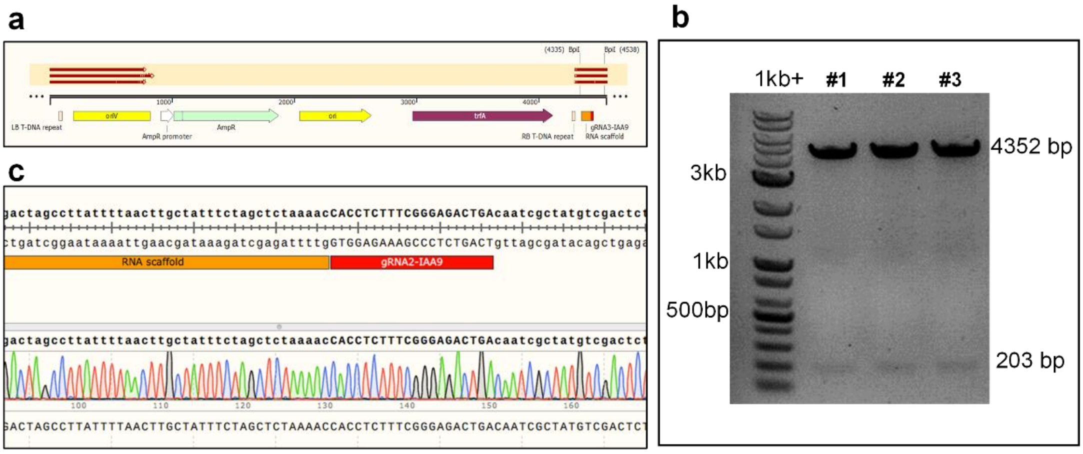


Supplementary Fig. 1 Cloning of *SlIAA-gRNA3* (level-1) construct. a Vector map and the positions of Sanger sequencing oligos in the plasmid; b *BpiI* digestion of the level-1 plasmids; c Sequencing result of clone #1.

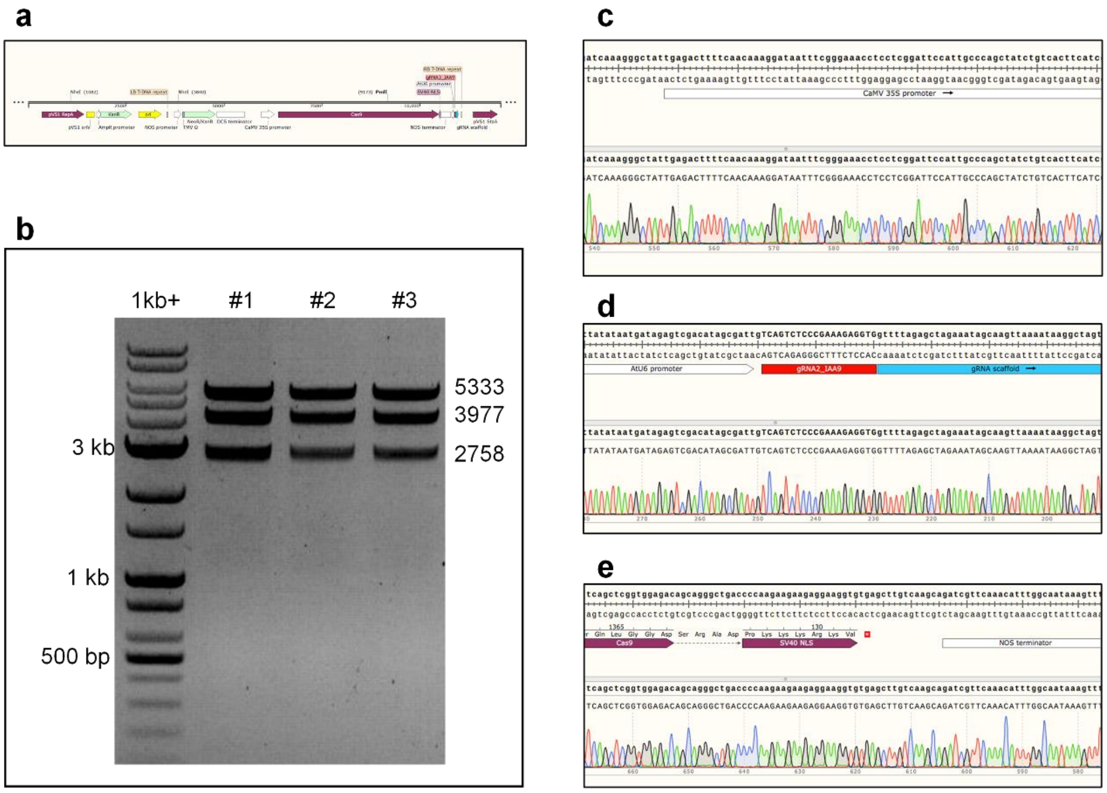


Supplementary Fig. 2 Cloning of *pGE-IAA9* (level-2) construct. a Vector map and the positions of Sanger sequencing oligos in the plasmid; b *NheI* and *PmlI* digestion of the level-2 plasmids; c-e Sequencing result of clone #1.

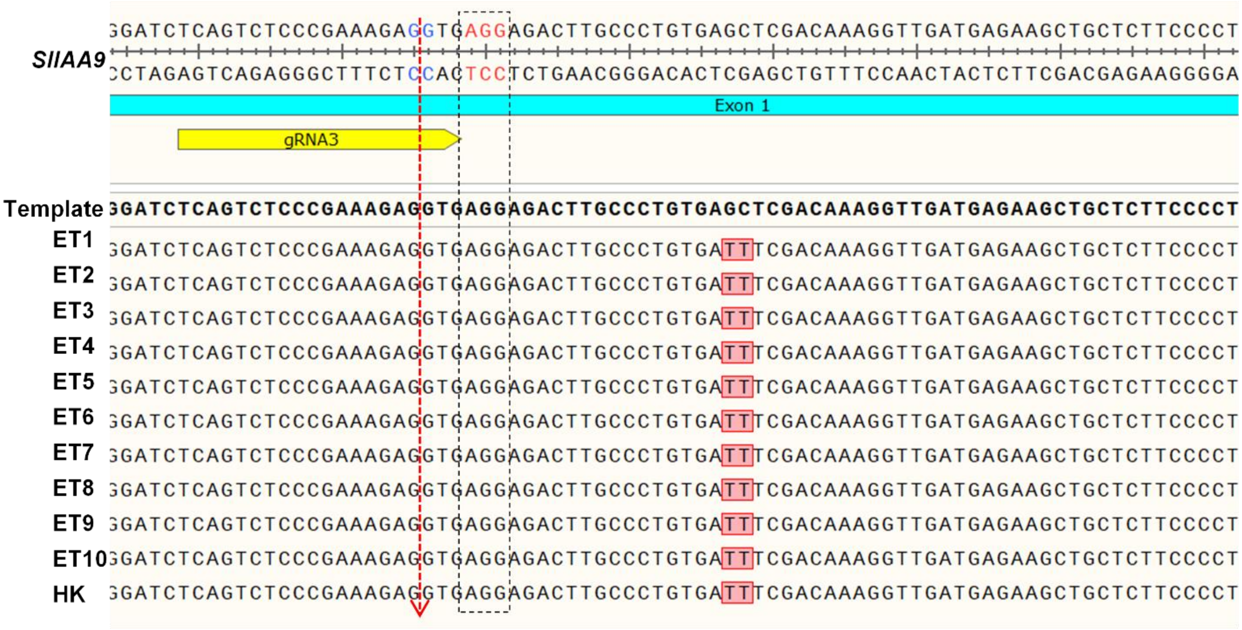


Supplementary Fig. 3 Sequences of editing frame in *SlIAA9* of ET lines. DNA samples were extracted from leave tissues of young leaves of ET lines and Hongkwang (HK), then analyzed by Sanger sequencing and Snapgene software. Template was extracted from sequencing data of *Solyc04g076850* from Sol Genomic Network (https://solgenomics.net/locus/463/view). Red dotted arrow indicated the Cas9 cleavage site in the target gene. The black box showed PAM sequence of the gRNA.

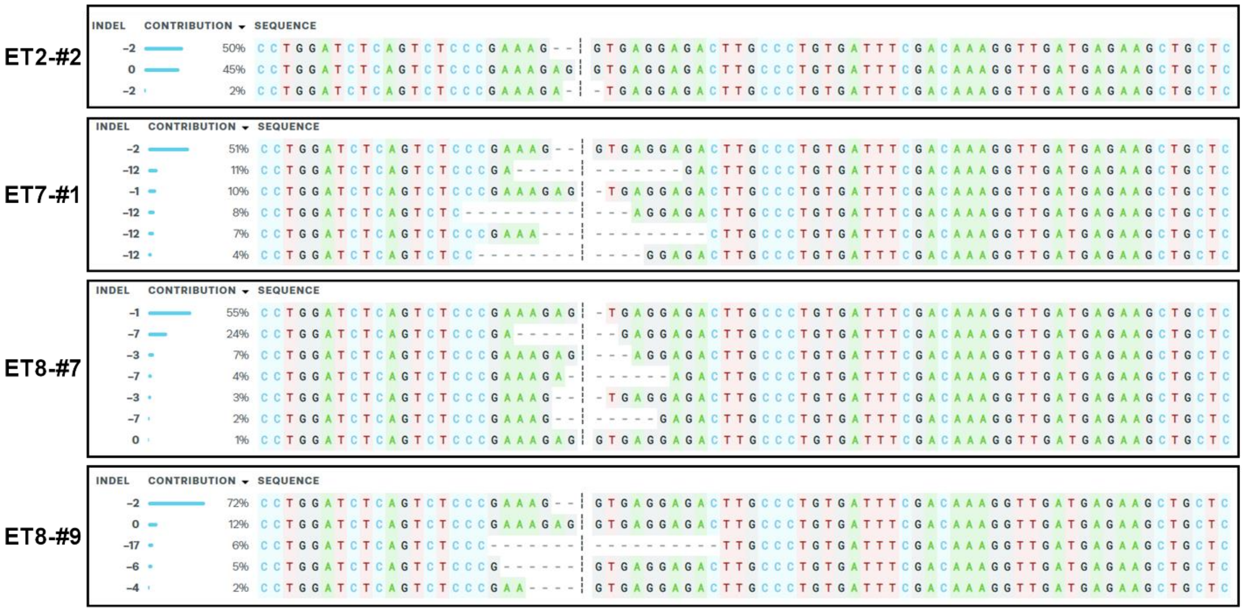


Supplementary Fig. 4 Allele contribution of selected G0 *SlIAA9*-edited plants.

Samples were taken from leaf tissues in three positions of each plant. DNA sequences were analyzed by Sanger sequencing and ICE Synthego software. These plants, together with ET5-#9 and ET8-#54, were selected to produce G1 by self-pollinating.
